# Longitudinal stability of individual brain plasticity patterns in blindness

**DOI:** 10.1101/2023.11.01.565196

**Authors:** Lénia Amaral, Peyton Thomas, Amir Amedi, Ella Striem-Amit

## Abstract

The primary visual cortex (V1) in individuals born blind is engaged in a wide spectrum of tasks and sensory modalities, including audition, touch, language, and memory. This widespread involvement raises questions regarding the constancy of its role and whether it might exhibit flexibility in its function over time, connecting to diverse network functions in response to task-specific demands. This would suggest that reorganized V1 takes on a role similar to cognitive multiple-demand system regions. Alternatively, it is possible that the varying patterns of plasticity observed in the blind V1 can be attributed to individual factors, whereby different blind individuals recruit V1 for different functions, highlighting the immense idiosyncrasy of plasticity. In support of this second account, we have recently shown that V1 functional connectivity varies greatly across blind individuals. But do these represent stable individual patterns of plasticity or merely instantaneous changes, for a multiple-demand system now inhabiting V1? Here we tested if individual connectivity patterns from the visual cortex of blind individuals are stable over time. We show that over two years, fMRI functional connectivity from the primary visual cortex is unique and highly stable in a small sample of repeatedly sampled congenitally blind individuals. Further, using multivoxel pattern analysis, we demonstrate that the unique reorganization patterns of these individuals allow decoding of participant identity. Together with recent evidence for substantial individual differences in visual cortex connectivity, this indicates there may be a consistent role for the visual cortex in blindness, which may differ for each individual. Further, it suggests that the variability in visual reorganization in blindness across individuals could be used to seek stable neuromarkers for sight rehabilitation and assistive approaches.

## Introduction

Complete sensory deprivation from birth offers the highest degree of plasticity attainable within the human brain, thereby inspiring extensive research endeavors focused on elucidating the developmental potential of a deprived cortex in the case of congenital blindness. Despite this long duration, the functional role of the early visual cortex in early-onset blindness remains unclear, with studies showing its recruitment for many different cognitive and sensory tasks, including audition, touch, smell, memory, language, and executive function^1-7^. This mixed pattern of findings, suggesting no single role for plastically reorganized early visual cortex (EVC), can be explained by two main hypotheses. It has been proposed that it plastically partake in many networks and functions depending on task demands over time, as part of a multiple-demand system^8^ which otherwise includes frontoparietal regions^9^. This would suggest a role in higher cognition and executive function for a typically early sensory system^8^, which would support high plasticity capacities for the human brain^3^ under such early-onset and extensive deprivation. While this suggestion has gained meaningful support, an alternative account, which is not completely mutually exclusive, exists. It is possible that at the group level, EVC seems to play many different roles, but in fact each blind individual utilizes their EVC for different functions, in a manner that is consistent over time. In support of this idea, we have recently shown that large variability exists among people with congenital blindness in how their visual cortex connects to other parts of the brain^10^. If this variability is consistent over time, this would support meaningful individual differences in the role of EVC due to blindness, which would reveal immense idiosyncrasy in plasticity and highlight the role of postnatal experience in forming individual brain organization patterns^10^.

Beyond the theoretical implications for our understanding of brain plasticity in humans, consistent individual differences in brain plasticity could also have translational value. Individual patterns of brain connectivity may be used to assess individuals’ risk, resilience, and even treatment efficacy for some disorders^11-15^, but this approach has not been attempted for rehabilitation in the case of blindness. Despite advances in medical restoration of visual information, through cataract removal^16-18^, gene therapy treatments^19,20^, stem-cell transplantation^21^, or retinal or cortical prostheses^22-27^, brain reorganization in blindness^3,28-34^ may limit the interpretation and processing of sight following early-onset or long-term blindness^35,36^. Accordingly, while some sight restoration is successful, large variability exists in functional outcome^37-40^, and it is hard to predict who will gain functional vision. If EVC has different connectivity and roles in each blind individual, this presents an opportunity to use individual patterns of functional connectivity (FC) as biomarkers for sight restoration efforts.

However, this requires that such profiles of connectivity be sufficiently stable over time to be relied on in prescribing long-term and potentially even invasive treatment. In the healthy, sighted, brain, Individual differences in connectivity appear to be stable across time over as much as 2.5 years^41-45^, suggesting they reflect true anatomical^46^ and functional differences, rather than merely different temporary cognitive states or motion patterns during the scans^44,47,48^. However, long-term stability in atypical (non-degenerative) brain reorganization has not been addressed. Specifically in blindness, if due to reorganization the EVC changes its connectivity and roles across time and tasks^8^, FC may only be able to provide a snapshot of an individual’s neural state at a particular point in time, and cannot be reliably used as a biomarker. Hence, it is crucial to validate the stability of FC in blindness to interpret the variability seen in blindness and potentially use it for individual fingerprinting.

Here, we tested FC profiles for consistency in individuals born blind over three scans performed across two years. These included both task-based fMRI scans and resting-state scans for these repeatedly-sampled individuals. We found that individual FC profiles were consistent over time and task performance. Further, consistent individual differences were found between each blind participant and a sighted cohort, suggesting stable individual patterns of visual cortex *plasticity* characterize each individual. Together, this provides evidence for the longitudinal stability of these measures in blindness, a persistent role for the early visual cortex in each blind individual, and their potential value as reliable biomarkers.

## Results

To inspect the stability of individual fingerprints of brain plasticity in blindness, we analyzed functional connectivity (FC) from eight repeatedly-sampled fully and congenitally blind individuals, which were taken over two years. These included three scanning sessions per participant: the first two included task data collected for studying the processing of auditory sensory substitution stimuli (published in ^49^), and were collected at an average of over one year apart (average time differences 15.5 months; see **Table 1** for participants characteristics and scan time differences). Each one of these sessions included two task runs. The third session collected resting-state data for the same participants, approximately a year after the previous session (^50^; on average 13.9 months between this and the previous scan; see **Fig. 1A**). To test for the stability of visual connectivity over time, we computed FC from an anatomically defined seed in retinotopic primary V1, based on a visual localizer in an independent group of sighted individuals^50^, while using task design as nuisance regressors for the task data (in addition to other nuisance regressors; see methods). Then, t-normalized V1-FC maps from all runs were analyzed for their consistency over time.

**Table 1.**
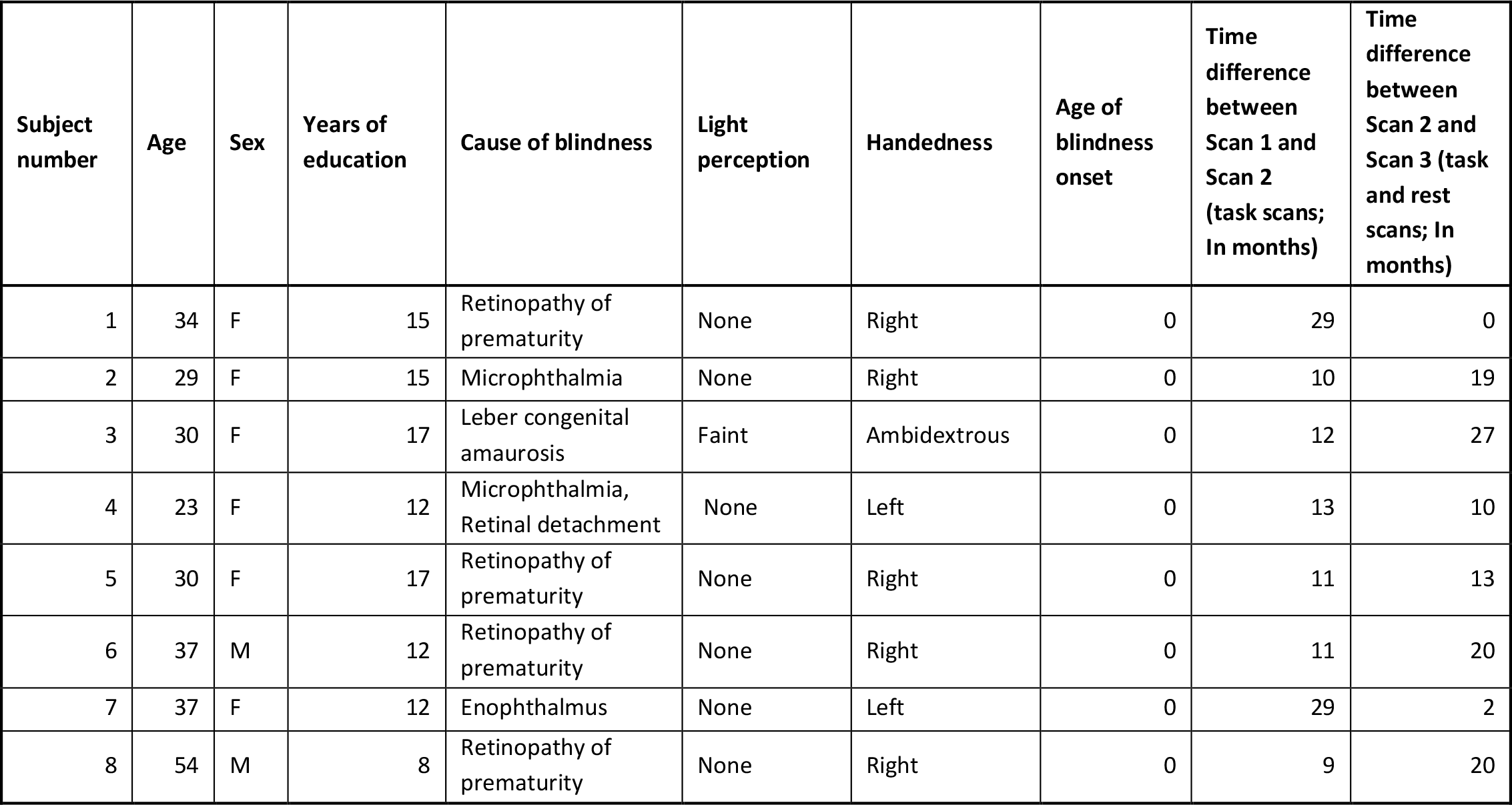
Characteristics of highly-sampled blind participants.

**Fig.1:**
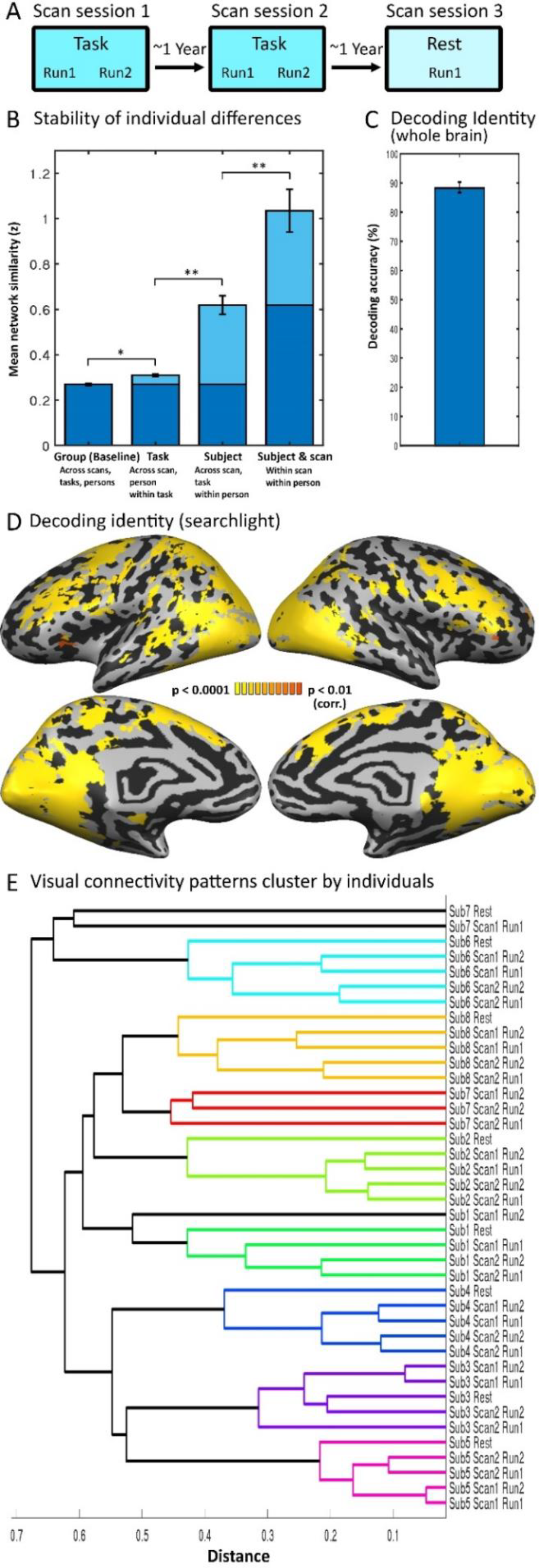
Stability of individual visual connectivity patterns in blindness A. Eight congenitally blind individuals were sampled in three scan sessions taken over two years: two task data sessions (2 runs each) and a resting-state session, scanned on average a year apart. These allowed us to test the stability of V1-FC over time in individuals born blind. B. The similarity of maps was tested by comparing the correlation between maps depending on shared features – across and within individuals, scans and tasks. Fisher-transformed correlation values are presented, showing the differential effect of task (task-based vs. resting-state scans), subject identity and same-day scans (only for task-data). * p<0.01 Bonf corr., ** p<0.001 Bonf. corr. C.Decoding for identity was calculated across scans for the whole-brain v1-FC data; overall decoding was highly significant (statistics calculated for the f1-factor, controlling for the unbalanced design; p<0.001). D.A decoding searchlight analysis for participant identity across time shows that vast areas of cortex, including in the visual ventral and dorsal pathways, along with dorsolateral and ventrolateral prefrontal cortex, had differential patterns of V1-FC for different blind individuals. E.A data-driven hierarchical clustering analysis of V1-FC maps for all five runs for each participant clustered primarily based on participant identity. Different color was assigned automatically to each group of nodes in the dendrogram whose linkage is less than 75%, yet marked subject identity largely accurately. This robustly shows the consistency of V1-FC maps over time within individuals. Sub-clustering was evident for task and scan effects. For the underlying similarity matrix see **Fig. S1**.

We used several approaches to test the stability of visual FC in blind individuals. (1) We performed correlations between the maps based on their participant identity and scan time. (2) We tested what factors contribute to the similarity of the maps by modeling different factors of similarity across the runs and testing their significance in explaining the variability of the maps. (3) We performed a cross-validated decoding analysis, testing if identity contributed to stable enough maps such that it can be assigned based on training on two time points to determine the identity at a different time point. (4) We performed a data-driven clustering analysis to qualitatively assess the similarity of the maps within each individual over time. (5) We tested the robustness of the individual similarity when adding data from additional blind and sighted individuals’ data. (6) Last, we tested if stable individual differences can be found in the plasticity patterns of the blind individuals, in the areas where their visual cortex is differently connected as compared to the sighted controls. Together, these analyses provided converging data supporting the consistency of individual V1-FC over time in blindness.

### Temporal stability of V1 connectivity patterns in blind individuals

First, we compared V1-FC maps for their similarity, using Pearson’s correlation, creating a similarity matrix across all 40 maps. To assess the contribution of within- and across-subject variables to the overall V1 FC network similarity, we averaged the similarity values for each scan condition independently from all others to evaluate task, subject, and scan type. We compared the average similarity values for each variable to the baseline similarity of all FC maps (**Fig. 1B**; Fisher-transformed values). Whether the scan was task or rest-based made a small but highly significant difference in the maps correlations (r=.30 vs r=0.26; t(446)=-5.59 p=1.83E-07, Bonf. corr.). However, subject identity contributed even more to the similarity of FC networks: V1-FC maps were significantly more similar for different time points for the same participant (r=0.53) than across participants (r=0.26; t(254)=16.67, p=3.42E-12, Bonf. corr.). Additionally, runs within a scan were more similar as compared to across scans for the same individual (r=0.73 vs 0.53; t(46)=4.09, p=1.72E-4 Bonf. corr.).

### Factors influencing V1 network similarity in blindness

To assess the explanatory value of individual identity in a data-driven manner, we computed a stepwise linear regression model to explore which factors explain the similarity matrix across the maps. This analysis added and removed variables to determine the most robust model (forward and backward stepwise regression). Input parameters included various explanatory factors, added as similarity models: overall similarity across all maps (null model, reflecting shared V1-FC for all the data), scan type (task/rest), session number (1-3), run number (1-2 for task data), participant identity (same/different, and if the scan was within the same session for the same individual. The data-driven resulting model included, beyond the overall similarity across all participants (the null model), variables of participant identity (coefficient estimate=0.33, t=21.2, p= 4.04E-79), task (coefficient estimate=0.09, t=4.45, p= 9.89E-06) and the within-session variable (same subject & scan; coefficient estimate=0.11, t=4.25, p=2.37E-05). Each of these factors were statistically significant and together accounted for 59.3% of the variability (R^2^= 0.593, F(775) vs. constant model=285, p = 1.04e-150). The addition of other parameters including run within session and session number did not prove statistically significant (each at p>0.05). Importantly, the introduction of head motion amplitude to the parameters used to generate the model, to control for similarity in the maps which could be driven by consistent head movement profiles for an individual across scans, did not change meaningfully the overall fit of the model (60.02% of the variability explained as compared to 59.3%). Although motion, the interaction between participant identity and task, and the interaction between task and motion were all significant (coefficient estimates=0.41, 0.06, 0.56, t=4.07, t=2.60, t=3.22, and p=5.19E-05, p=9.38E-03, p=1.31E-03 respectively), participant identity (coefficient estimate=0.33, t=21.4, p= 2.41E-80), task (coefficient estimate=1.21E-01, t=5.38, p= 9.89E-08), and the within-session variable (same subject & scan; coefficient estimate=0.11, t=4.19, p=3.14E-05) remained the factors with the most explanatory value, as in the original model. This suggests stable individual patterns of connectivity significantly contribute to the variability in the data.

### Using the stability of individual patterns to decode participant identity

To test if individual patterns are stable enough over time to allow decoding of participant identity, we trained a linear classifier on the distinction between participants’ data from two scanning sessions and tested it on data collected on a third scanning session, taken (on average) a year apart. Statistical testing was performed on the f1 value, which better accounts for unbalanced false negative and false positive errors^51^, given that in this case the chance level of a true positive would be 1/8, but the chance level of a true negative would be 7/8. Decoding of participant identity was highly significant as compared to a permutation test (**Fig. 1C**; accuracy = 88 ± 1.9%, F1-score = 51 ± 12%, precision = 57 ± 9.7%, recall = 53 ± 17.5%, chance level=12.5%, p<0.0001); in fact, decoding of each subject’s data compared to the others was significant (each p<0.05, vs an independent permutation tests), showing stable unique patterns of visual connectivity. To examine where in the brain individual patterns of V1-FC differ enough for allowing decoding, we performed a whole-brain searchlight decoding analysis. Functional connectivity patterns from V1 to vast areas in the bilateral ventral and dorsal visual streams contributed to decoding identity, as well as dorsolateral, ventrolateral frontal cortex and the superior frontal gyrus (**Fig. 1D**). Connectivity to auditory cortex and early sensorimotor cortex did not contain as much discernable identity information. This suggests that connectivity from V1 to vast areas of cortex is stable enough over time to allow identifying individuals.

### Qualitative assessment of the individual consistency of V1 connectivity

To visually explore the similarity across the runs, we used a hierarchical clustering approach on subjects’ V1-FC whole-brain (gray-matter masked) maps. Distance was calculated as the correlation between individual FC map data. A dendrogram of the distances across all participants was computed based on the weighted distance between clusters; different color was assigned automatically to each group of nodes in the dendrogram whose linkage is less than 75%. This data-driven approach yielded robust results (**Fig. 1E;** see dissimilarity matrix in **Fig.S1**): all but three maps (37/40, 92%) were colored by participant identity, showing a clear clustering by stable identity patterns. As a sub-clustering, the data did further cluster by scan type and scan session; showing that some of the variability does stem from cognitive state (task performance as compared to resting) and potentially temporal factors unique for each scan day.

### Idiosyncrasy of V1 network patterns among a larger cohort

The previous analyses provide converging data supporting the stability of individual differences in blind adults. Nevertheless, does this clustering stem from the limited quantity of data, or do blind individuals have unique stable patterns when considering a larger population of participants? To test this, we similarly analyzed V1-FC maps from seventeen additional congenitally blind participants, each of whom participated in a single resting-state run^50,52^. Of these, five were scanned in the same scanner and parameters as the resting-state data from our highly-sampled cohort, and twelve were scanned elsewhere^52^. Despite the addition of more plastically-modified patterns of visual connectivity, scans of our highly-sampled participants continued to cluster together for each individual (again, 37/40, 92%; **Fig. S2A**). Last, we similarly analyzed V1-FC maps from seventeen control (sighted) participants scanned during rest in the same scanning conditions as the highly-sampled individuals (overall matching the blind cohort in age and education^50,52^). Again, scans of our highly-sampled blind participants clustered together (37/40, 92%; **Fig. S2B**), and show higher similarity within as compared to across participants than across participants (r=0.53 vs r=0.23; t(254)=14.07, p=3.72E-33, Bonf. corr; see **Fig. S2C**). A full network similarity analysis including these additional sighted controls showed that task (rest vs. task data; t(1982)=10.43, p=1.6E-24 Bonf. corr.) and subject identity both contributed a significant amount to the similarity of FC, with subject contributing the largest effect (**Fig. S2C**). Group (blind/sighted) contributed a significant but modest effect (t(3739)=3.45, p=0.0006 Bonf. corr.).

### Long-term stability in visual cortex plasticity patterns

Overall, our data supported high stability of individual patterns of visual connectivity in blindness. Our recent work showed that blind individuals differ from one another more in areas whose connectivity is affected by blindness^10^. Are the plasticity patterns in each blind person, i.e. changes to FC as compared to the typical sighted system, also stable over time? To test this, we first tested if brain areas of stable individual differences correspond to areas that show brain plasticity in blindness. We contrasted V1-FC between our sample of repeatedly-sampled individuals and the control group to show which areas show plasticity in blindness at the group level (**Fig. S3A**). We then calculated the correlation between the areas showing changes in RSFC at the group level (**Fig. S3A**) and the decoding map of participant identity across time (**Fig. 1D**). The two maps were highly correlated (CCC=0.439; p<0.0001; **Fig. S3B** for permutation analysis); this suggests that changes in V1-FC in blindness form individual stable features.

To examine if plasticity patterns are stable more directly, we then computed the difference between each V1-FC map of the highly-sampled individuals and the group of sighed controls scanned at the same scan site using the same scan parameters (using a Crawford modified t-test^58,59^, appropriate for case study statistics). This provided us with maps showing the unique pattern of changes between each such scan of the blind participant and the corresponding control group. Importantly, these plasticity maps were also consistent within participants, showing similar trends as the V1-FC maps; The correlation within participant was higher than across participants (**Fig. 2A**; r=0.38 vs r=0.10, t(254)=14.71, p=2.34E-35, Bonf. corr.), identity decoding was similarly high and significant (**Fig. 2B**; f1=52%±11.9; accuracy = 89%±1.8; precision = 58 ± 9.0%; recall = 55 ± 17.9%, all significant at p<0.0001), and these maps similarly clustered by participant identity (**Fig. 2C**). Together, these results show that individual patterns of visual cortex plasticity are consistent over time in individuals born blind.

**Fig.2:**
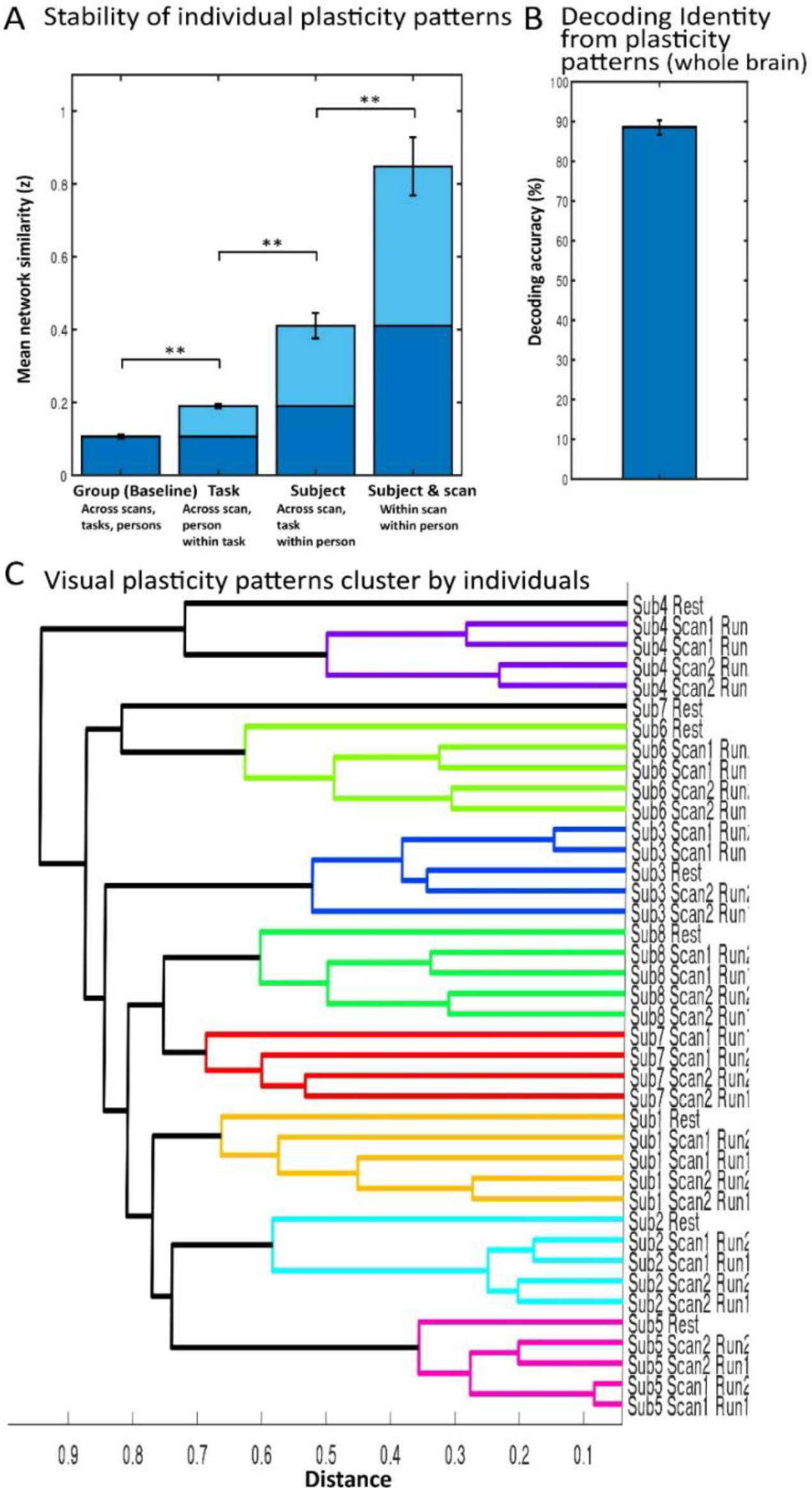
Individual patterns of V1 connectivity plasticity are stable over time in blindness Analyses presented in Fig. 1 were repeated for maps of changes in V1-FC for each run of the repeatedly sampled blind individuals as compared to V1-FC from a control sighted cohort (Crawford t-test). These analyses reflect the stability of plasticity patterns of V1-FC in each individual. A. The similarity of V1-FC maps was tested by comparing the correlation between maps depending on shared features – across and within individuals, scans and tasks. Fisher-transformed correlation values are presented, showing the differential effect of task (task-based vs. resting-state scans), subject identity and same-day scans (only for task-data). Plasticity patterns showed high correlation within participant as compared to across participants. ^*^ p<0.01 Bonf corr., ^**^ p<0.0001 Bonf. corr. B.Decoding accuracy for identity from individual plasticity maps was calculated across scans; overall decoding was highly significant (statistical testing was based on the f1 value, accounting for the unbalanced design; p<0.001). C.A data-driven hierarchical clustering analysis of V1-FC plasticity maps for all five runs for each participant clustered primarily based on participant identity. Different color was assigned automatically to each group of nodes in the dendrogram whose linkage is less than 75%, yet marked subject identity largely accurately. This robustly shows the consistency of plasticity patterns of V1-FC over time within individuals. Sub-clustering was evident for task and scan effects.

## Discussion

The functional role of the Early Visual Cortex (EVC) in congenital blindness remains unclear, given its recruitment for many sensory and cognitive tasks. This complex response pattern alludes to flexible use of deprived cortex over time, whereby the blind brain may allocate the connectivity and use of the visual cortex based on task demands^8^. Here we show that this is not the case: in the context of blindness, subject identity plays a notably more substantial role in explaining the variability in V1 FC as compared to the nature of the task and to the passing time. Individual patterns of visual cortex connectivity are extremely consistent over a period of two years (**Fig. 1**), allowing decoding of one’s identity from their brain connectivity pattern (**Fig. 1C**,**D**). Additionally, consistency is found in the pattern of plasticity (**Fig. 2**) – the aspect in which each blind participant’s brain differs from the sighted control group, suggesting plasticity, though variable *across* individuals^10^, is highly stable over time and task *within* individuals in blindness. This suggests that the blind brain may not be flexible over time in its use of the visual cortex across task demands^8^, but rather has a persistent connectivity pattern stable over years, across cognitive states as different as an highly-demanding task and rest. We have recently shown that large individual differences in visual cortex connectivity exist in blindness^10^. The stability of these individual patterns also suggests that neural fingerprinting could be used for individually-tailored rehabilitation from blindness. These theoretical and translational aspects will be discussed in turn in the next sections.

First, the stability of individual differences of FC across task and rest has been validated in typically developed sighted controls^44,47,48^, showing an underlying effect of individual brain pattern across states. The stability of brain-wide FC profiles has also been validated in typically-developed individuals over as much as 2.5 years^41-43^. However, in congenital blindness there is additional reason for concern as to the stability of visual cortex connectivity patterns; the role of the early visual cortex, the site of possible rehabilitation interventions, remains highly debated^3,33^. Over the decades since neuroimaging allowed the inspection of visual cortex in blindness, the primary visual cortex in congenital blindness has been shown to respond to a vast variety of tasks, from language and memory to tactile and auditory perception, including echolocation and braille reading^1,2,4,6,7,60-63^. This complex pattern of group-level results lead to speculation that the early visual cortex in blindness is tuned through attention, and may be form a multiple-demand cortex that can flexibly adjust its connectivity depending on task demands^8^. This, in turn, suggested similarity between V1 function in blindness and higher-level cortical areas in the brain, for example, the frontal lobe^3,64^. In our results, V1-FC to the frontal cortex is variable enough across individuals on one hand, and consistent enough within individuals over time on the other hand, to allow for decoding of identity over the course of two years (**Fig. 1D**). More broadly, in showing the large individual differences found across blind individuals in the connectivity of the primary visual cortex^10^, and in showing their overall stability over time, we propose an alternative account: that group level results showing the involvement of V1 in multiple tasks stemmed from individuals whose visual cortex has plastically organized differently, and that maintains its plastically-formed connectivity over time in adulthood.

We cannot, at this point, determine the origin of these individual differences, and they likely reflect a combination of differential inherited predisposition^65,66^ for the connectivity of EVC, as well as the effects of early-onset and lifelong use of compensatory senses, cognitive faculties and strategies^2,4,67-77^Similarly, although future research should test when and how these individual differences arise developmentally, by adulthood, a blind individual appears to have a stable pattern of connectivity for their primary visual cortex. To clarify, we do not claim there is nothing in common to changes in EVC’s connectivity across blind individuals, or question the importance of group-level studies in blindness. Many consistent effects of plasticity are highly reproducible across studies and cohorts around the world, clearly showing cross-modal plasticity in blind EVC can be found for many functions (e.g. different responses for language across modalities (^1,7,61,78,79^ though possibly also in sighted people^80^). Importantly, several studies in the past have also acknowledged the large variability existing in the observed plasticity in blindness (e.g. ^79,81-83^, including for language ^84^). Some have even shown that this variability has functional significance, showing correlation between the variable behavioral compensatory capacities in blindness and the level of EVC recruitment for different tasks (e.g.^2,4,77^). In our data as well, a common pattern of FC exists across blind participants (denoted as “group” in **Figs. 1B** and **2B**), accounts for a significant part of the variability in our data. However, at least in this sample of highly-sampled individuals, individual differences explain a larger portion of the variance of the different scans.

Likewise, we do not wish to argue that state-based changes are impossible in the blind visual cortex; indeed, we find significant effects of task on the FC of the visual cortex, and see significantly higher correlation between brain patterns obtained within the same day and task than across them (**Fig. 1B**; at least for the auditory task, as we only collected one resting-state run). Our data does not allow us to adjudicate if these stem from technical issues such as scanner calibration or the participant’s temporary state. Optimally, future studies including many types of tasks and resting-state within the same scan day in blindness would assess this question. However, our data suggests that individual patterns of V1 connectivity plasticity are overall very stable over time and states – across an engaging auditory task and resting-state. This is especially noteworthy given the relative short duration of each of the scans involved (^∼^10 minutes each) as compared to some deep-data studies focusing on individuals^85-87^, as individual functional connectivity measures tend to stabilize further with longer acquisition periods^88-92^(though meaningful stability is already achieved in 10 minutes^92^). However, even with relatively shorter scans, stability is clear, suggesting this is possibly an underestimation of these effects.

This stability, in turn, could have implications for using individual patterns of brain plasticity for tailoring assistive and rehabilitative solutions based on each individual’s brain pattern, similarly to the attempt for neuromarkers for other conditions^11-15^. This may assist in determining the likelihood a blind individual would benefit from any of a large number of approaches available to them. Given the dizzying selection of sensory substitution techniques^93-98^ and sensory aids (e.g. for reading: refreshable Braille displays, screen readers, and optical and electronic aids using touch, audition, and vision, respectively; in navigation even more options exist^99,100^), narrowing these down to fit the individual would be valuable. This is even more important in applying invasive medical restoration approaches^101^ (e.g. cataract removal^16-18^, gene therapy treatments^19,20^, stem-cell transplantation^21^, or retinal or cortical prostheses^22-27,102^), given the high variability in outcomes in these cases^37-40^; despite invasive procedures, some patients gain little functional sight. Since some of this variability may stem from neural causes, if the visual system is reorganized in a manner that does not allow restoration of the original function ^35^, the search for neuromarkers is apt. Future research would be needed to show the link between these stable brain patterns and behavior, and further, their usefulness for rehabilitation. However, here we take the first crucial step for using them for prospective treatment, by establishing that these patterns, and particularly plasticity patterns for each blind individual, are stable in adulthood.

## Methods

### Participants

Eight congenitally-blind individuals participated in the study. The data was collected for two previous studies^49,50^, scanned at three separate dates. Participants were between 23 and 54 years of age at the time of the last scan. See **Table 1** for detailed characteristics of the blind participants and scan time differences. The Tel-Aviv Sourasky Medical Center Ethics Committee approved the experimental procedure. Written informed consent was obtained from each participant.

### Functional Imaging

Functional magnetic resonance imaging (fMRI) data were obtained during two experiments which included sensory substitution task performance and resting conditions, without any external stimulation or task (e.g., spontaneous blood oxygen level-dependent fluctuations) for each participant. During the resting state scan, subjects lay supine in the scanner with no external stimulation or explicit task. During the task scans, subjects heard sounds conveying visual information as translated through sensory substitution^94^ and performed location (left vs right location) and shape (round vs angular shapes) judgements^49^. Specifically, in a slow event related design, during each trial, subjects were presented with an auditory instruction: either “shape” or “location”, which directed their attention to the task. They were then presented with a 1-s soundscape (sensory substitution sound rendering of the visual stimulus) that was repeated 4 times (total presentation time—4 s), given 5 additional seconds to reconstruct the image in their mind and were then instructed, by an auditory cue, to respond using a response box. Subjects used a two-button response box to indicate the parameter specified (Is the shape round? Is it in the left of the picture?). The full detail of the experimental design are described in ^49^.

Images were acquired with a 3-T General Electric scanner with an InVivo 8-channel head coil. Task data was comprised of two 315 whole-brain functional volumes per scan date, acquired with an Echo Planar Imaging sequence (Repetition Time = 1500 ms, Echo Time = 30 ms, 29 slices, voxel size 3 × 3 × 4 mm, flip angle 70°, scan duration =7.9 minutes). Resting-state data was comprised of one functional run, containing 180 continuous whole-brain functional volumes acquired with an Echo Planar Imaging sequence (Repetition Time = 3000 ms, Echo Time = 30 ms, 29-46 slices, voxel size 3 × 3 × 4 mm, flip angle 90°, 182 volumes, scan duration = 9.1 min). T1-weighted anatomical images were acquired using a 3D MPRAGE sequence (typical scan parameters were: 58 slices; TR = 8.9 ms; TE = 3.5 ms; inversion time = 450ms; FA = 13°; FOV = 256 × 256 mm; voxel size = 1 × 1 × 1 mm; matrix size = 256 × 256).

### fMRI preprocessing

Data analysis was performed using the BrainVoyager 20 software package (Brain Innovation, Maastricht, Netherlands) and custom scripts in MATLAB (MathWorks, Natick, MA) following standard preprocessing procedures. The first two images of each scan were excluded due to non-steady state magnetization. Preprocessing of functional scans included 3-Dimensional motion correction, slice scan time correction and band pass filtering (0.01-0.1Hz). Head motion did not exceed 2 mm along any given axis or include spike-like motion of more than 1 mm in any direction. Functional and anatomical data sets for each subject were aligned and fit to standardized Talairach space^103^.

### Seed regions-of-interest

The region-of-interest (ROI) for the primary visual cortex (V1) was defined from an independent localizer, acquired in a separate group of 14 sighted subjects^50^ using a standard phase-encoded retinotopic mapping protocol, with eccentricity and polar mapping of ring and wedge stimuli, respectively^104-107^. The full experimental detail can be found in ^50^. Polar mapping data were used to define the borders of V1, used as a seed ROI for the FC analyses.

### FC analyses

Individual time courses from the V1 seed ROI were sampled from each of the participants’ runs, z-transformed and used as individual predictors in a z-normalized GLM analysis. Individual motion estimates (6 degrees of freedom and their first derivatives) and signals from the ventricles and white matter regions (defined using the grow-region function in Brain Voyager on the individual level) were used as nuisance predictors. For task data, to inspect the underlying functional connectivity pattern beyond the effects of the task, nuisance predictors also included the experimental conditions’ predictors, including the different tasks (shape, location), and separating periods of stimulus presentation and response. Individual FC maps were masked with a gray matter mask. To inspect individual variability only in areas which may play a role in connectivity to visual cortex in blindness, analyses of similarity were performed within areas showing a significant FC to V1 at the blind group level (group level random-effect GLM of all runs used from the highly sampled individuals, at p<0.05 corrected for multiple comparisons using the spatial extent method, a set-level statistical inference correction; ^108,109^).

### Additional participants

V1-RSFC data from seventeen additional congenitally blind individuals and seventeen sighted controls, published in ^10^ was included in the analysis to test the stability of individual differences when compared with additional exemplars of V1-FC. The data were collected for two previous studies ^50,52^; only resting state data is analyzed here. During the resting state scan, subjects lay supine in the scanner with no external stimulation or explicit task. Sighted participants kept their eyes closed. The data for the blind participants was scanned at two separate sites. Data for five of the additional blind participants and all seventeen sighted participants used here was collected at the same location and scanning parameters as the repeatedly-sampled blind individuals used for the main analyses (Cohort A; ^50^). Sighted participants had normal or corrected-to-normal vision, all participants had no history of neurologic disorder, and were matched to the repeatedly-sampled blind group in age (t(24)=1.38, p=0.18) and education (t(24)=-0.90, p=0.38). The Tel-Aviv Sourasky Medical Center Ethics Committee approved the experimental procedure. Data for the additional twelve blind participants was collected at a different scanner (Cohort B; see full parameters in ^52^), and the Institutional Review Board of the Department of Psychology, Peking University, China and the Institutional Review Board of Harvard University approved the experimental procedure. **Table S1** provides detailed characteristics of the additional blind participants whose data was included. In all cases, written informed consent was obtained from each participant. Data from these additional participants was analyzed similarly to the main dataset, generating V1-FC maps.

### Network similarity analysis

To evaluate the contribution of each scan variable on the overall similarity matrix of the V1-FC networks (Fischer transformed), the average similarity contribution was calculated for the effects of task condition (task/rest), subject identity, and scan session. Group similarity was used as the baseline comparison, and was calculated from the average similarity across all unique FC values, from different subjects, tasks, and scan sessions (n=224). Task similarity contribution was then averaged from all unique task FC values (same task, different subject, and different scan, n=224). Subject was similarly calculated from all values with the same subject, different task, and different scan (n=32). Within-subject and within-scan variables were calculated from all FC values of the same subject, same scan session, but different task run (n=16); these were calculated only for task data (sessions 1, 2) as the session in which resting-state data was acquired did not include additional runs available for this analysis. Significant differences in variable similarity contributions were calculated from a t-test, Bonferroni corrected for multiple comparisons. The network similarity analysis was similarly conducted on FC dissimilarity matrices derived from other blind and sighted participants (**Fig.S2C)**. In this case, average group datapoints (n=1760) were used as the baseline to compare the additional similarity contributions of task (n=224) and subject (n=32).

To test whether head motion is similar within each participant over time, and has a significant role in the similarity of the networks over time in blindness, we used head movement as an additional model for the network similarity analysis. We computed the Fourier-transformed amplitude of movement frequency in each run. This was done while adjusting for the different duration of the runs, by using the shared duration of the data (157 TRs). The amplitude of movement in each direction was averaged and used to create a similarity matrix, which was inputted to the network similarity analysis.

### Regression models

Following the calculation of the dissimilarity matrix between runs, the correlation values of lower non-diagonal elements of the dissimilarity matrix were statistically compared between subjects on different runs of the same date, across dates and across task and rest scans. This was done both as simple Spearman correlation (**Fig. 1B**) and in fitting a stepwise linear regression model to account for the relative contribution of each of the variables to the explained variance.

### Decoding analysis

To test if individual patterns of FC were stable over time in blind individuals, we performed a multivariate multivoxel pattern analysis using CosMoMVPA^110^. We used a linear discrimination analysis (LDA) classifier to discriminate between t-normalized V1-FC patterns of the 8 participants across different scan time points using a cross-validation scheme. For example, the classifier was trained using data from scan time point 1 and 2, and tested using data from scan time point 3. This cross-validation ensured that training and testing data was kept completely independent and enable us to access the capacity of the classifier to generalize across different time points. Finally, we calculated the following decoding metrics for the group average: accuracy, precision, recall and f1-score. The F1-score is a balanced measure that considers both false positives and false negatives, providing a single value to evaluate the model’s performance^51^. Since our false positives and false negatives did not have a similar cost (i.e., the chance level of a true positive would be 1/8, but the chance level of a true negative would be 7/8), F1-score was more appropriate for significance testing. The group F1-score was statistically tested using non-parametric Monte Carlo sampling by comparing the probability of each true F1-scores to its group level null-distribution with 10,000 null accuracies created by randomly shuffling the condition labels (i.e., subject identity). Single-subject permutation tests were also computed to assess the decoding ability for each individual blind brain pattern (10,000 permutations). In order to inspect where in the brain the decoding occurs, 100 null maps were generated for each participant, by randomly shuffling the condition labels ^111^. These chance-F1 maps were then sampled with 10,000 iterations to generate a null distribution, which was used to generate group level statistics, corrected for multiple comparisons as above, at the cluster level (spatial extent method; ^108,109^). The resulting p-values were converted to z-values as implemented in CoSMoMVPA.

### Clustering analysis

To test the consistency of individual patterns of FC from the visual cortex, we performed a hierarchical clustering analysis across subjects V1-seeded FC maps from across different scans. Distance was calculated as the correlation between individual FC vectors, implemented in MATLAB (MathWorks, Natick, MA). A dendrogram of the distances across all participants was computed based on complete distance between clusters (**Fig. 1E**; for the underlying correlation dissimilarity matrix see **Fig. S1**). Different color was assigned automatically to each group of nodes in the dendrogram whose linkage is less than 75%. The hierarchical clustering was also similarly conducted on individual maps derived from various other blind and sighted participants (**Fig. S2A**,**B**; there different coloring schemes thresholds were used), and for the plasticity stability analysis (see below; **Fig. 2**).

### Plasticity stability

to assess if the unique patterns of changes between each blind participant’s brain and the sighted typical brain is also stable over time, we calculated the Crawford modified t-test^58,59^ between each V1-FC map and the V1-FC of sighted control group participants. These maps reflect the plasticity of the V1-FC pattern of each run of the blind participant. These maps were then analyzed using comparable network similarity, decoding, and clustering analyses as those performed on the V1-FC maps (**Fig. 2**; see above).

## Acknowledgements

We are thankful to the blind subjects who participated in our experiment. We also thank Maria Czarnecka for her comments on this work. This work was supported by the NIH NEI/OBSSR grant no. 1R01EY034515 and the Edwin H. Richard and Elisabeth Richard von Matsch Distinguished Professorship in Neurological Diseases (to ESA).

## Notes

### Competing Interest Statement

The authors have declared no competing interest.

## References

1 Burton, H., Diamond, J. B. & McDermott, K. B. Dissociating cortical regions activated by semantic and phonological tasks: a FMRI study in blind and sighted people. J Neurophysiol 90, 1965-1982. Epub 2003 Jun 1964. (2003).

2 Amedi, A., Raz, N., Pianka, P., Malach, R. & Zohary, E. Early ‘visual’ cortex activation correlates with superior verbal memory performance in the blind. Nat Neurosci 6, 758–766. (2003).

3 Bedny, M. Evidence from Blindness for a Cognitively Pluripotent Cortex. Trends in Cognitive Sciences 21, 637-648, doi:10.1016/j.tics.2017.06.003 (2017).

4 Gougoux, F., Zatorre, R. J., Lassonde, M., Voss, P. & Lepore, F. A functional neuroimaging study of sound localization: visual cortex activity predicts performance in early-blind individuals. PLoS Biol 3, e27 (2005).

5 Weeks, R. et al. A positron emission tomographic study of auditory localization in the congenitally blind. J Neurosci 20, 2664–2672. (2000).

6 Stilla, R. et al. Neural processing underlying tactile microspatial discrimination in the blind: a functional magnetic resonance imaging study. J Vis 8, 13 11-19, doi:10.1167/8.10.13 (2008).

7 Sadato, N. et al. Activation of the primary visual cortex by Braille reading in blind subjects. Nature 380, 526–528. (1996).

8 Pelland, M. et al. State-dependent modulation of functional connectivity in early blind individuals. NeuroImage 147, 532–541, doi:10.1016/j.neuroimage.2016.12.053 (2017).

9 Duncan, J. The multiple-demand (MD) system of the primate brain: mental programs for intelligent behaviour. Trends in Cognitive Sciences 14, 172–179, doi:10.1016/j.tics.2010.01.004 (2010).

10 Sen, S. et al. The Role of Visual Experience in Individual Differences of Brain Connectivity. J Neurosci 42, 5070-5084, doi:10.1523/JNEUROSCI.1700-21.2022 (2022).

11 Foulkes, L. & Blakemore, S.-J. Studying individual differences in human adolescent brain development. Nature Neuroscience 21, 315-323, doi:10.1038/s41593-018-0078-4 (2018).

12 Rosenberg, M. D. et al. A neuromarker of sustained attention from whole-brain functional connectivity. Nature Neuroscience 19, 165, doi:10.1038/nn.4179 (2015).

13 Friedman, N. P. & Miyake, A. Unity and diversity of executive functions: Individual differences as a window on cognitive structure. Cortex 86, 186–204, doi:10.1016/j.cortex.2016.04.023 (2017).

14 Drysdale, A. T. et al. Resting-state connectivity biomarkers define neurophysiological subtypes of depression. Nature Medicine 23, 28, doi:10.1038/nm.4246 (2016).

15 Fox, M. D. & Greicius, M. Clinical applications of resting state functional connectivity. Frontiers in systems neuroscience 4, 19-19, doi:10.3389/fnsys.2010.00019 (2010).

16 Ostrovsky, Y., Andalman, A. & Sinha, P. Vision following extended congenital blindness. Psychol Sci 17, 1009–1014 (2006).

17 Maurer, D. Critical periods re-examined: Evidence from children treated for dense cataracts. Cognitive Development, doi:10.1016/j.cogdev.2017.02.006 (2017).

18 Kalia, A. et al. Assessing the impact of a program for late surgical intervention in early-blind children. Public Health 146, 15–23, doi:10.1016/j.puhe.2016.12.036 (2017).

19 Ashtari, M. et al. Plasticity of the human visual system after retinal gene therapy in patients with Leber’s congenital amaurosis. Science Translational Medicine 7, 296ra110, doi:10.1126/scitranslmed.aaa8791 (2015).

20 Sahel, J. A. & Roska, B. Gene therapy for blindness. Annu Rev Neurosci 36, 467-488, doi:10.1146/annurev-neuro-062012-170304 (2013).

21 Singh, M. S. et al. Reversal of end-stage retinal degeneration and restoration of visual function by photoreceptor transplantation. Proc Natl Acad Sci U S A 110, 1101-1106, doi:10.1073/pnas.1119416110 (2013).

22 Beauchamp, M. S. et al. Dynamic Stimulation of Visual Cortex Produces Form Vision in Sighted and Blind Humans. Cell 181, 774-783.e775, doi:10.1016/j.cell.2020.04.033 (2020).

23 Bosking, W. H., Beauchamp, M. S. & Yoshor, D. Electrical Stimulation of Visual Cortex: Relevance for the Development of Visual Cortical Prosthetics. Annual Review of Vision Science 3, 141-166, doi:10.1146/annurev-vision-111815-114525 (2017).

24 Luo, Y. H.-L. & da Cruz, L. The Argus® II Retinal Prosthesis System. Progress in Retinal and Eye Research, doi:10.1016/j.preteyeres.2015.09.003.

25 Weiland, J. D., Liu, W. & Humayun, M. S. Retinal prosthesis. Annual Review of Biomedical Engineering 7, 361-401, doi:10.1146/annurev.bioeng.7.060804.100435 (2005).

26 Roux, S. et al. Probing the functional impact of sub-retinal prosthesis. eLife 5, e12687. doi:10.7554/eLife.12687 (2016).

27 Maya-Vetencourt, J. F. et al. A fully organic retinal prosthesis restores vision in a rat model of degenerative blindness. Nature materials, doi:10.1038/nmat4874 (2017).

28 Frasnelli, J., Collignon, O., Voss, P. & Lepore, F. Crossmodal plasticity in sensory loss. Prog Brain Res 191, 233-249, doi:10.1016/B978-0-444-53752-2.00002-3 (2011).

29 Voss, P. & Zatorre, R. J. Organization and reorganization of sensory-deprived cortex. Curr Biol 22, R168-173, doi:10.1016/j.cub.2012.01.030 (2012).

30 Kupers, R. & Ptito, M. Compensatory plasticity and cross-modal reorganization following early visual deprivation. Neurosci Biobehav Rev, doi:10.1016/j.neubiorev.2013.08.001 (2013).

31 Heimler, B., Striem-Amit, E. & Amedi, A. Origins of task-specific sensory-independent organization in the visual and auditory brain: neuroscience evidence, open questions and clinical implications. Curr Opin Neurobiol 35, 169-177, doi:10.1016/j.conb.2015.09.001 (2015).

32 Cecchetti, L., Kupers, R., Ptito, M., Pietrini, P. & Ricciardi, E. Are Supramodality and Cross-Modal Plasticity the Yin and Yang of Brain Development? From Blindness to Rehabilitation. Frontiers in Systems Neuroscience 10, doi:10.3389/fnsys.2016.00089 (2016).

33 Fine, I. & Park, J.-M. Blindness and Human Brain Plasticity. Annual Review of Vision Science 4, 337-356, doi:10.1146/annurev-vision-102016-061241 (2018).

34 Bridge, H. & Watkins, K. E. Structural and functional brain reorganisation due to blindness: The special case of bilateral congenital anophthalmia. Neuroscience & Biobehavioral Reviews 107, 765–774, doi:10.1016/j.neubiorev.2019.10.006 (2019).

35 Striem-Amit, E., Bubic, A. & Amedi, A. in Frontiers in the Neural Bases of Multisensory Processes (eds M.M. Murray & M. T. Wallace) 393–420 (Taylor and Francis, 2011).

36 Kiorpes, L. The Puzzle of Visual Development: Behavior and Neural Limits. The Journal of Neuroscience 36, 11384-11393, doi:10.1523/jneurosci.2937-16.2016 (2016).

37 Carlson, S., Hyvarinen, L. & Raninen, A. Persistent behavioural blindness after early visual deprivation and active visual rehabilitation: a case report. Br J Ophthalmol 70, 607–611 (1986).

38 Ganesh, S. et al. Results of late surgical intervention in children with early-onset bilateral cataracts. British Journal of Ophthalmology 98, 1424-1428, doi:10.1136/bjophthalmol-2013-304475 (2014).

39 Gregory, R. L. & Wallace, J. G. in Experimental Psychology Society, Monograph Supplement. 2 (Heffers, 1963).

40 Huber, E. et al. A Lack of Experience-Dependent Plasticity After More Than a Decade of Recovered Sight. Psychological Science, doi:10.1177/0956797614563957 (2015).

41 Jovicich, J. et al. Longitudinal reproducibility of default-mode network connectivity in healthy elderly participants: A multicentric resting-state fMRI study. Neuroimage 124, 442-454, doi:10.1016/j.neuroimage.2015.07.010 (2016).

42 Badhwar, A. et al. Multivariate consistency of resting-state fMRI connectivity maps acquired on a single individual over 2.5 years, 13 sites and 3 vendors. Neuroimage 205, 116210, doi:10.1016/j.neuroimage.2019.116210 (2020).

43 Liu, W. et al. Longitudinal test-retest neuroimaging data from healthy young adults in southwest China. Scientific Data 4, 170017, doi:10.1038/sdata.2017.17 (2017).

44 Gratton, C. et al. Functional Brain Networks Are Dominated by Stable Group and Individual Factors, Not Cognitive or Daily Variation. Neuron 98, 439-452.e435, doi:10.1016/j.neuron.2018.03.035 (2018).

45 Arazi, A., Gonen—Yaacovi, G. & Dinstein, I. The magnitude of trial-by-trial neural variability is reproducible over time and across tasks in humans. eneuro, doi:10.1523/eneuro.0292-17.2017 (2017).

46 Honey, C. J. et al. Predicting human resting-state functional connectivity from structural connectivity. Proc Natl Acad Sci U S A 106, 2035-2040, doi:10.1073/pnas.0811168106 (2009).

47 Finn, E. S. et al. Functional connectome fingerprinting: identifying individuals using patterns of brain connectivity. Nature neuroscience (2015).

48 Tavor, I. et al. Task-free MRI predicts individual differences in brain activity during task performance. Science 352, 216-220, doi:10.1126/science.aad8127 (2016).

49 Striem-Amit, E., Dakwar, O., Reich, L. & Amedi, A. The large-scale organization of “visual” streams emerges without visual experience. Cereb Cortex 22, 1698-1709, doi:10.1093/cercor/bhr253 (2012).

50 Striem-Amit, E. et al. Functional connectivity of visual cortex in the blind follows retinotopic organization principles. Brain 138, 1679-1695, doi:10.1093/brain/awv083 (2015).

51 Chinchor, N. & Sundheim, B. M. in Fifth Message Understanding Conference (MUC-5): Proceedings of a Conference Held in Baltimore, Maryland, August 25-27, 1993.

52 Striem-Amit, E., Wang, X., Bi, Y. & Caramazza, A. Neural representation of visual concepts in people born blind. Nat Commun 9, 5250, doi:10.1038/s41467-018-07574-3 (2018).

53 Liu, Y. et al. Whole brain functional connectivity in the early blind. Brain 130, 2085–2096 (2007).

54 Yu, C. et al. Altered functional connectivity of primary visual cortex in early blindness. Hum Brain Mapp 29, 533–543 (2008).

55 Wang, D. et al. Altered resting-state network connectivity in congenital blind. Human Brain Mapping 35, 2573-2581, doi:10.1002/hbm.22350 (2013).

56 Burton, H., Snyder, A. Z. & Raichle, M. E. Resting state functional connectivity in early blind humans. Front Syst Neurosci 8, 51, doi:10.3389/fnsys.2014.00051 (2014).

57 Qin, W., Xuan, Y., Liu, Y., Jiang, T. & Yu, C. Functional Connectivity Density in Congenitally and Late Blind Subjects. Cerebral Cortex, doi:10.1093/cercor/bhu051 (2014).

58 Crawford, J. R. & Garthwaite, P. H. Comparison of a single case to a control or normative sample in neuropsychology: Development of a Bayesian approach. Cognitive Neuropsychology 24, 343-372, doi:10.1080/02643290701290146 (2007).

59 Crawford, J. R. & Howell, D. C. Comparing an Individual’s Test Score Against Norms Derived from Small Samples. The Clinical Neuropsychologist 12, 482-486, doi:10.1076/clin.12.4.482.7241 (1998).

60 Burton, H., Snyder, A. Z., Diamond, J. B. & Raichle, M. E. Adaptive changes in early and late blind: a FMRI study of verb generation to heard nouns. J Neurophysiol 88, 3359–3371. (2002).

61 Lane, C., Kanjlia, S., Omaki, A. & Bedny, M. “Visual” Cortex of Congenitally Blind Adults Responds to Syntactic Movement. The Journal of Neuroscience 35, 12859-12868, doi:10.1523/jneurosci.1256-15.2015 (2015).

62 Abboud, S. & Cohen, L. Distinctive Interaction Between Cognitive Networks and the Visual Cortex in Early Blind Individuals. Cereb Cortex 29, 4725-4742, doi:10.1093/cercor/bhz006 (2019).

63 Thaler, L., Arnott, S. R. & Goodale, M. A. Neural correlates of natural human echolocation in early and late blind echolocation experts. PLoS ONE 6, e20162. doi:10.1371/journal.pone.0020162 (2011).

64 Deen, B., Saxe, R. & Bedny, M. Occipital Cortex of Blind Individuals Is Functionally Coupled with Executive Control Areas of Frontal Cortex. Journal of Cognitive Neuroscience, 1-15, doi:10.1162/jocn_a_00807 (2015).

65 Molloy, M. F. & Saygin, Z. M. Individual Variability in Functional Organization of the Neonatal Brain. NeuroImage, 2021.2003.2024.436788, doi:10.1016/j.neuroimage.2022.119101 (2022).

66 Ortiz-Terán, L. et al. Brain circuit–gene expression relationships and neuroplasticity of multisensory cortices in blind children. Proceedings of the National Academy of Sciences, doi:10.1073/pnas.1619121114 (2017).

67 Röder, B. et al. Improved auditory spatial tuning in blind humans. Nature 400, 162–166 (1999).

68 Van Boven, R. W., Hamilton, R. H., Kauffman, T., Keenan, J. P. & Pascual-Leone, A. Tactile spatial resolution in blind braille readers. Neurology 54, 2230–2236 (2000).

69 Goldreich, D. & Kanics, I. M. Tactile acuity is enhanced in blindness. J. Neurosci. 23, 3439–3445 (2003).

70 Beaulieu-Lefebvre, M., Schneider, F. C., Kupers, R. & Ptito, M. Odor perception and odor awareness in congenital blindness. Brain Res Bull 84, 206-209, doi:10.1016/j.brainresbull.2010.12.014 (2011).

71 Tillman, M. H. & Bashaw, W. L. Multivariate analysis of the WISC scales for blind and sighted children. Psychol Rep 23, 523–526 (1968).

72 Pozar, L. Effect of long-term sensory deprivation on recall of verbal material. Studia Psychologica 24, 311–311 (1982).

73 Raz, N., Striem, E., Pundak, G., Orlov, T. & Zohary, E. Superior Serial Memory in the Blind: A Case of Cognitive Compensatory Adjustment. Curr Biol 17, 1129-1133, doi:10.1016/j.cub.2007.05.060 (2007).

74 Occelli, V., Lacey, S., Stephens, C. & Sathian, K. Superior verbal abilities in congenital blindness. Electronic Imaging 2016, 1-4, doi:10.2352/ISSN.2470-1173.2016.16.HVEI-094 (2016).

75 Dormal, V., Crollen, V., Baumans, C., Lepore, F. & Collignon, O. Early but not late blindness leads to enhanced arithmetic and working memory abilities. Cortex, doi:10.1016/j.cortex.2016.07.016 (2017).

76 Loiotile, R., Omaki, A. & Bedny, M. Enhanced performance on a sentence comprehension task in congenitally blind adults. Language, Cognition and Neuroscience (2019).

77 Voss, P. & Zatorre, R. J. Occipital Cortical Thickness Predicts Performance on Pitch and Musical Tasks in Blind Individuals. Cereb Cortex, doi:10.1093/cercor/bhr311 (2011).

78 Bedny, M., Pascual-Leone, A., Dodell-Feder, D., Fedorenko, E. & Saxe, R. Language processing in the occipital cortex of congenitally blind adults. Proc Natl Acad Sci U S A 108, 4429-4434, doi:10.1073/pnas.1014818108 (2011).

79 Abboud, S., Engemann, D.-A. & Cohen, L. Semantic coding in the occipital cortex of early blind individuals. bioRxiv, 539437, doi:10.1101/539437 (2019).

80 Seydell-Greenwald, A., Wang, X., Newport, E. L., Bi, Y. & Striem-Amit, E. Spoken language processing activates the primary visual cortex. PLoS One 18, e0289671. doi:10.1371/journal.pone.0289671 (2023).

81 van den Hurk, J., Van Baelen, M. & Op de Beeck, H. P. Development of visual category selectivity in ventral visual cortex does not require visual experience. Proceedings of the National Academy of Sciences 114, E4501-E4510, doi:10.1073/pnas.1612862114 (2017).

82 Mattioni, S. et al. Categorical representation from sound and sight in the ventral occipitotemporal cortex of sighted and blind. eLife 9, e50732. doi:10.7554/eLife.50732 (2020).

83 Rosenke, M. et al. Extensive individual differences of category information in ventral temporal cortex in the congenitally blind. bioRxiv, 2020.2006.2014.151092, doi:10.1101/2020.06.14.151092 (2020).

84 Watkins, K. E., Coullon, G. S. L. & Bridge, H. Language and nonverbal auditory processing in the occipital cortex of individuals who are congenitally blind due to anophthalmia. Neuropsychologia 173, 108304, doi:10.1016/j.neuropsychologia.2022.108304 (2022).

85 Gratton, C., Nelson, S. M. & Gordon, E. M. Brain-behavior correlations: Two paths toward reliability. Neuron 110, 1446-1449, doi:10.1016/j.neuron.2022.04.018 (2022).

86 Nee, D. E. fMRI replicability depends upon sufficient individual-level data. Communications Biology 2, 130, doi:10.1038/s42003-019-0378-6 (2019).

87 Chen, G. et al. Hyperbolic trade-off: The importance of balancing trial and subject sample sizes in neuroimaging. NeuroImage 247, 118786, doi:10.1016/j.neuroimage.2021.118786 (2022).

88 Laumann, Timothy O. et al. Functional System and Areal Organization of a Highly Sampled Individual Human Brain. Neuron 87, 657–670, doi:10.1016/j.neuron.2015.06.037 (2015).

89 Finn, E. S. et al. Can brain state be manipulated to emphasize individual differences in functional connectivity? NeuroImage 160, 140–151, doi:10.1016/j.neuroimage.2017.03.064 (2017).

90 Birn, R. M. et al. The effect of scan length on the reliability of resting-state fMRI connectivity estimates. NeuroImage 83, 550–558, doi:10.1016/j.neuroimage.2013.05.099 (2013).

91 Mueller, S. et al. Reliability correction for functional connectivity: Theory and implementation. Human Brain Mapping 36, 4664–4680, doi:10.1002/hbm.22947 (2015).

92 Airan, R. D. et al. Factors affecting characterization and localization of interindividual differences in functional connectivity using MRI. Human Brain Mapping 37, 1986–1997, doi:10.1002/hbm.23150 (2016).

93 Bach-y-Rita, P. Tactile sensory substitution studies. Ann N Y Acad Sci 1013, 83–91 (2004).

94 Meijer, P. B. An experimental system for auditory image representations. IEEE Trans Biomed Eng 39, 112–121 (1992).

95 Capelle, C., Trullemans, C., Arno, P. & Veraart, C. A real-time experimental prototype for enhancement of vision rehabilitation using auditory substitution. IEEE Trans Biomed Eng 45, 1279-1293, doi:10.1109/10.720206 (1998).

96 Striem-Amit, E., Guendelman, M. & Amedi, A. ‘Visual’ acuity of the congenitally blind using visual-to-auditory sensory substitution. PLoS One 7, e33136. doi:10.1371/journal.pone.0033136 (2012).

97 Martolini, C. et al. in 2018 IEEE International Symposium on Medical Measurements and Applications (MeMeA). 1–6.

98 Maimon, A. et al. The Topo-Speech sensory substitution system as a method of conveying spatial information to the blind and vision impaired. Frontiers in Human Neuroscience 16, doi:10.3389/fnhum.2022.1058093 (2023).

99 Kuriakose, B., Shrestha, R. & Sandnes, F. E. Tools and Technologies for Blind and Visually Impaired Navigation Support: A Review. IETE Technical Review 39, 3-18, doi:10.1080/02564602.2020.1819893 (2022).

100 Mashiata, M. et al. Towards assisting visually impaired individuals: A review on current status and future prospects. Biosensors and Bioelectronics: X 12, 100265, doi:10.1016/j.biosx.2022.100265 (2022).

101 Ptito, M. et al. Brain-Machine Interfaces to Assist the Blind. Frontiers in Human Neuroscience 15, doi:10.3389/fnhum.2021.638887 (2021).

102 Bosking, W. H. et al. Percepts evoked by multi-electrode stimulation of human visual cortex. Brain Stimulation: Basic, Translational, and Clinical Research in Neuromodulation 15, 1163-1177, doi:10.1016/j.brs.2022.08.007 (2022).

103 Talairach, J. & Tournoux, P. Co-planar stereotaxic atlas of the human brain. (Thieme, 1988).

104 Engel, S. A. et al. fMRI of human visual cortex. Nature 369, 525, doi:10.1038/369525a0 (1994).

105 Sereno, M. I. et al. Borders of multiple visual areas in humans revealed by functional magnetic resonance imaging. Science 268, 889–893 (1995).

106 Wandell, B. A., Dumoulin, S. O. & Brewer, A. A. Visual field maps in human cortex. Neuron 56, 366-383, doi:10.1016/j.neuron.2007.10.012[doi] (2007).

107 Wandell, B. A. & Winawer, J. Imaging retinotopic maps in the human brain. Vision Research 51, 718-737, doi:10.1016/j.visres.2010.08.004 (2011).

108 Forman, S. D. et al. Improved assessment of significant activation in functional magnetic resonance imaging (fMRI): use of a cluster-size threshold. Magn Reson Med 33, 636–647 (1995).

109 Friston, K. J., Worsley, K. J., Frackowiak, R. S. J., Mazziotta, J. C. & Evans, A. C. Assessing the significance of focal activations using their spatial extent. Human Brain Mapping 1, 210-220, doi:10.1002/hbm.460010306 (1993).

110 Oosterhof, N. N., Connolly, A. C. & Haxby, J. V. CoSMoMVPA: multi-modal multivariate pattern analysis of neuroimaging data in Matlab / GNU Octave. Frontiers in Neuroinformatics 10, 27, doi:10.1101/047118 (2016).

111 Stelzer, J., Chen, Y. & Turner, R. Statistical inference and multiple testing correction in classification-based multi-voxel pattern analysis (MVPA): Random permutations and cluster size control. NeuroImage 65, 69–82, doi:10.1016/j.neuroimage.2012.09.063 (2013).

