## Supplemental Material for "Longitudinal stability of individual brain plasticity patterns in blindness"

**Longitudinal stability of individual brain plasticity patterns in blindness – Supplemental material**

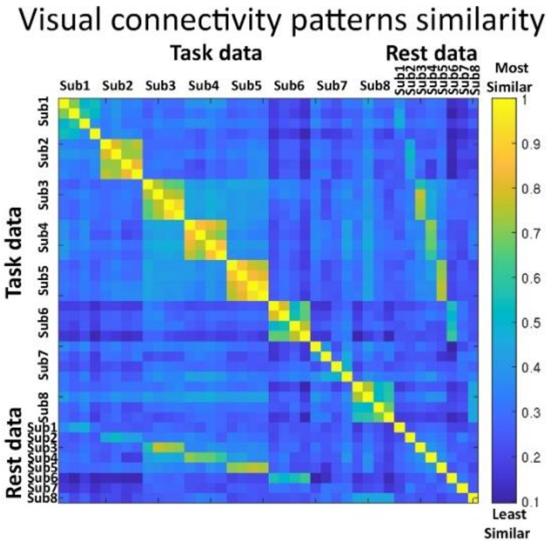


Fig. S1:
The V1-RSFC correlation (similarity) structure between individuals based on which hierarchical clustering analysis (Fig. 1E) was conducted.

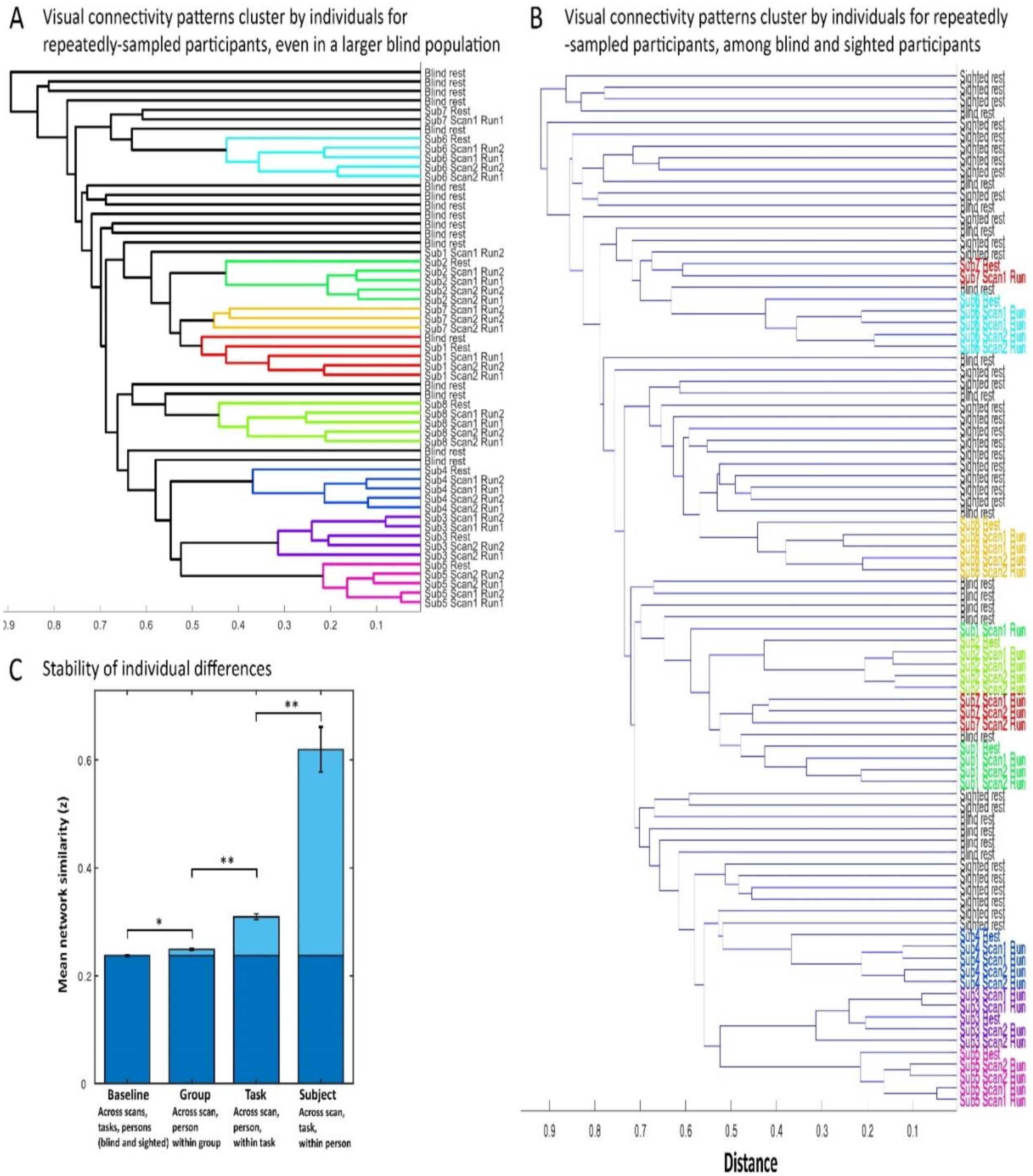


Fig.S2: Individual visual connectivity patterns in blindness are consistent and unique even when compared to a variety of blind and sighted data.

**A.** The data-driven hierarchical clustering analysis of V1-FC maps (**Fig. 1E**) was repeated with the addition of V1-RSFC maps from seventeen additional congenitally blind participants (listed as “Blind rest” in the dendrogram; one map per participant). Despite the larger possibility for mixing of maps for the repeatedly-sampled individuals, their V1-Fc maps from across five runs over three years continued to cluster together. Different color was assigned automatically to each group of nodes in the dendrogram whose linkage is less than 45%, robustly showing the consistency of V1-FC maps over time within individuals. Sub-clustering was evident for task and scan effects.

**B.** The data-driven hierarchical clustering analysis of V1-FC maps (**panel A**) was repeated with the addition of V1-RSFC maps from seventeen sighted participants (listed as “Sighted rest” in the dendrogram; one map per participant). Despite the larger possibility for mixing of maps for the repeatedly-sampled individuals, their V1-Fc maps from across five runs over three years continued to cluster together for the most part. Colors of participant names are manually added to demonstrate the consistent clustering based on individual identity among the many additional patterns of V1-FC.

**C.** The similarity of V1-FC maps for all blind and sighted participants (8 repeatedly-sampled blind participants, 17 additional blind participants and 17 control participants) was tested by comparing the correlation between maps depending on shared features – across and within group individuals, scans and tasks. Fisher-transformed correlation values are presented, showing the differential effect of group (blind vs. sighted), task (task-based vs. resting-state scans), subject identity and same-day scans (only for task-data). * $p<0.01$ Bonf corr., ** $p<0.0001$ Bonf. corr.

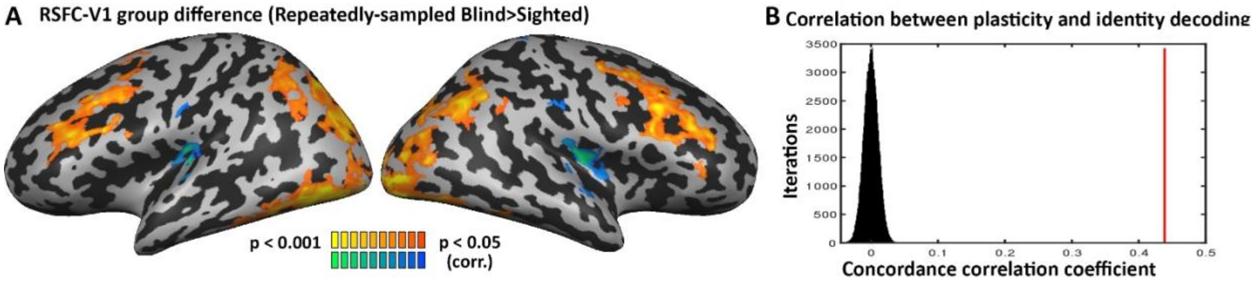


Fig. S3: Areas that show V1-FC plasticity in blindness include unique individual patterns stable over time

**A.** The difference in V1-FC between the eight repeatedly-sampled blind participants and the control group (n=17) is depicted. As reported before[50,53-57], the blind and sighted differed in their RSFC from the primary visual cortex to visual, parietal, and frontal regions.

**B.** Overall across the brain, areas showing changes in RSFC in the eight highly-sampled individuals (**panel A**) corresponded to areas where participant decoding was significant across time (**Fig. 1D**). The concordance correlation coefficient was calculated between the RSFC group difference and decoding maps (red line; CCC=0.439) and compared with a spatial permutation test (distribution in black; $p<0.0001$).

| Subject number | Age | Sex | Years of education | Cause of blindness | Light perception | Handedness | Age of blindness onset | Cohort |
|---|---|---|---|---|---|---|---|---|
| 7 | 23 | M | 12 | Microphthalmia | None | Right | 0 | A |
| 9 | 31 | M | 12 | Retinopathy of prematurity | None | Right | 0 | A |
| 10 | 35 | F | 12 | Retinoblastoma | None | Right | 0 | A |
| 11 | 34 | F | 17 | Microphthalmia | None | Left | 0 | A |
| 13 | 42 | M | 17 | Retinopathy of prematurity | Faint | Right | 0 | A |
| 14 | 36 | M | 12 | Microphthalmia | None | Ambidextrous | 0 | B |
| 15 | 22 | M | 15 | Microphthalmia | None | Right | 0 | B |
| 16 | 33 | M | 12 | Microphthalmia; microcornea | None | Right | 0 | B |
| 17 | 48 | M | 12 | Glaucoma | None | Right | 0 | B |
| 18 | 46 | F | 9 | Glaucoma | None | Right | 0 | B |
| 19 | 40 | M | 12 | Leukoma | Faint | Right | 0 | B |
| 20 | 50 | F | 12 | Cataracts; eyeball dysplasia | Faint | Right | 0 | B |
| 21 | 57 | M | 12 | Eyeball dysplasia | None | Right | 0 | B |
| 22 | 43 | F | 12 | Glaucoma | None | Right | 0 | B |
| 23 | 48 | M | 12 | Microphthalmia; cataracts; leukoma | None | Right | 0 | B |
| 24 | 63 | M | 9 | Glaucoma; leukoma | None | Right | 0 | B |
| 25 | 41 | F | 12 | Optic nerve atrophy | Faint | Right | 0 | B |

Table S1: Characteristics of additional blind participants.

Cohort A was acquired in Israel and comprised of additional 5 blind adults[50]. Cohort B was acquired in China and comprised of 12 blind adults[52].
